# CFTR dysfunction in the intestinal epithelium is sufficient to promote pathogenic expansion of *E. coli* and enhanced barrier permeability in cystic fibrosis

**DOI:** 10.64898/2026.09.18.752678

**Authors:** Philip P. Ahern, Naseer Sangwan, Apollo Stacy, Jessica M. Snyder, Zaria Johnson, Megan Cua, Brian Chang, Alexander Maynard, Lauren Smith, Filip Sagl, Brandon Bakos, Akeem Santos, Tina Nunn, Anne-Marie C. Overstreet, Michael Wannemuehler, Samuel I. Miller, Jeannette S. Messer, Mohammed Dwidar, Mitchell L. Drumm, Craig Hodges, Adeline M. Hajjar

## Abstract

Changes in the gut microbiome in cystic fibrosis (CF) are well characterized, yet their causes and downstream effects remain largely unknown. Well-documented alterations include reduced overall complexity (i.e. alpha diversity) of the gut microbiota and increased relative abundance of *E. coli*, which are associated with greater inflammation and shorter stature in infants.

Previous results from our laboratory using a germ-free cystic fibrosis transmembrane conductance regulator (*Cftr*) mutant mouse model (CF mouse) demonstrated that the observed fecal microbiome dysbiosis is driven by mutated *Cftr* independent of factors such as diet or antibiotic treatment. We expand on these results in this report by using the defined 8-member community Altered Schaedler Flora (ASF) with and without *E. coli*, to show that *E. coli* is pathogenic in the context of the CF gut microbiome, resulting in increased intestinal permeability. We also show that *Cftr* deletion in intestinal epithelial cells alone, using a Villin-Cre targeted model, is sufficient to raise *E. coli* abundance in the fecal microbiome, increase intestinal permeability, and amplify the number of TH17 cells in the mesenteric lymph nodes.

Together, our results demonstrate that the intestinal epithelium plays a dominant role in fecal microbiome alterations in CF and that the resultant dysbiosis contributes to CF pathogenesis.

## 2. Introduction

Cystic fibrosis (CF) is a genetic disorder caused by mutations in the cystic fibrosis transmembrane conductance regulator (*Cftr*) gene, leading to dysfunction of the CFTR chloride channel and resulting in dehydrated mucus [1]. CF is associated with multisystem complications, most notably involving the respiratory and gastrointestinal (GI) tracts. Common GI complications include intestinal inflammation and malabsorption, even after treatment of pancreatic insufficiency [2]. A key physiological manifestation of GI disease in CF is impaired mucosal barrier function, observed in people with CF more than 30 years ago [2–4]. Mouse CF models also show significantly impaired mucosal barrier function compared to control mice [5], especially in the small intestine, where tight junctions were observed to be disrupted [6]. The distal small intestine also serves as a site of bacterial overgrowth [7,8] and intestinal obstruction [9,10], both amplified by slower gut motility [11,12]. Structural changes include abnormal Paneth cell granule accumulation in intestinal crypts, reducing antimicrobial function [13].

CF is strongly linked to gut microbiota dysbiosis [14]. Overall lower alpha diversity in the gut microbiota has been characterized in stool samples from children with CF compared to healthy controls [15,16]. Elevated levels of pro-inflammatory bacteria, specifically proteobacteria (now called pseudomonadota) such as *Escherichia coli* (*E. coli*), have been characterized in the altered CF microbiome in children with CF, and correlate with higher fecal calprotectin as well as shorter stature [15,17–19]. The microbial environment of the CF gut may impose selective pressures that drive rapid parallel genetic adaptations in commensal *E. coli* that enhance its survival in the inflamed CF gut [20]. Further supporting the pathogenic role of gut microbiome dysbiosis, CFTR modulator therapy, used to improve CFTR function, has been linked to decreased intestinal inflammation and reduced *E. coli* abundance [21].

Using a gnotobiotic mouse model, we previously showed that the CF-specific dysbiotic gut microbial community is driven by CF genotype (i.e. *Cftr* mutation) and not by factors like diet and antibiotics common in human CF treatments [22,23]. Furthermore, the CF-specific microbiota was maintained only in a CF mouse, as the microbiome reverted to a control-like community when transplanted into control non-CF mice. CFTR dysfunction therefore drives the selection of the gut microbiome, leading to an increased relative abundance of *E. coli* and elevated inflammatory markers compared to control mice [22]. Additionally, decreasing the bacterial load in CF mice by using antibiotics decreased inflammatory markers [24], suggesting microbiome dysregulation plays a dominant role in GI inflammation.

Here, we use a reductionist approach. We colonized germ-free CF and control mice with a defined 8-member bacterial community, the Altered Schaedler Flora (ASF) [25], with or without *E. coli*, to test whether we can replicate *E. coli* expansion in the CF gut. ASF was developed in the 1970s as a community of bacteria isolated from mice that is stable across multiple generations, and that lacks colonization resistance [26,27]. We used this defined community to determine whether *E. coli* expansion in CF can be attributed to intestinal barrier dysfunction. In addition, we previously showed that crypts in the distal small intestine of germ-free CF mice were dilated, likely from dehydrated mucus accumulation, with entrapment of Paneth cell granules independent of the microbiota [22]. Therefore, to test the role of *Cftr* expression in intestinal epithelial cells, we rederived the mouse line with intestinal-epithelium targeted *Cftr* deletion (Villin-Cre mediated deletion of a floxed *Cftr* allele [28]) to germ-free status, isolating the role of intestinal epithelium-specific loss of *Cftr* from that of other cell types (e.g., immune cells) when colonized with ASF and *E. coli*.

## 3. Methods

### 3.1 Mice

The germ-free *Cftr*^tm1Unc^ (S489X, [29]) CF line has been previously described [22]. Mice are bred through heterozygous crosses and homozygous knockout (KO) mice are referred to as CF mice, whereas non-CF mice include both wild-type and heterozygous mice. The *Cftr*^*fl*/fl^ x Villin-Cre line [28] was rederived germ-free via cesarean section at the Cleveland Clinic Gnotobiotics Facility. The intestinal KO group consists of *Cftr*^*fl*/fl^ Vil:Cre^+^ mice, whereas intestinal controls are a combination of *Cftr*^*fl*/fl^ Vil:Cre^-^, *Cftr*^*fl*/+^ Vil:Cre^+^, or *Cftr*^*fl*/+^ Vil:Cre^-^. All gnotobiotic mice were maintained on cellulose paper bedding (AlphaDri® PLUS, Shepherd Specialty Papers) with an autoclaved chow diet (LabDiet 5021) and autoclaved non-acidified Milli-Q water. All experimental mice (controls and knockouts) were maintained on laxative water as previously described [22], to prevent intestinal obstructions [30]. Mice were housed in a facility with a 14hr light and 10hr dark cycle. Mice were not randomized because we matched the age and sex of mice between KO and control groups of each line. Specific pathogen-free (SPF) *Cftr*^*G542X*^ (G542X, [31]) mice, which are nearly identical in phenotype to the S489X mouse line, and wild-type littermates used in Figure 1A, were housed in standard polysulfone microisolator cages in ventilated racks with corncob bedding. SPF mice were fed irradiated Rodent Diet 7904 (Teklad Diets Envigo, Madison WI) ad libitum and were provided a laxative in acidified (pH 2.5-3) and autoclaved water in water bottles and maintained on a 12hr light and 12hr dark cycle.

**Figure 1:**
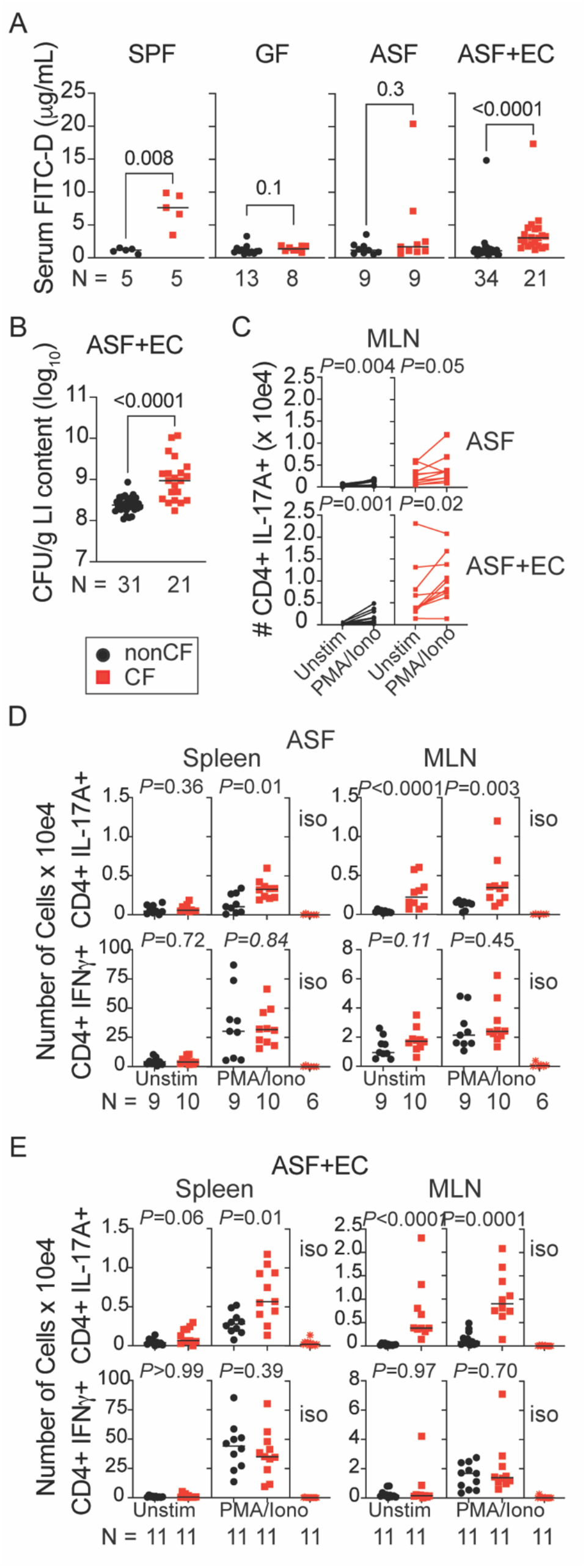
ASF + *E. coli* disrupts intestinal barrier in CF mice with increased LI CFU and TH17 cells in spleen and MLN. **Panel A)** FITC-dextran was measured in the serum 4 hr after oral administration to overnight fasted mice in conventional mice (SPF), germ-free mice (GF), ASF-colonized (ASF) and ASF + *E. coli* (ASF+EC; 7-37 d post colonization with *E. coli*) colonized mice. **Panel B)** Colon contents (LI, large intestine) collected at necropsy of Panel A mice were plated on MacConkey II Agar for *E. coli* CFU enumeration. **Panels C, D & E)** Cells from spleens or mesenteric lymph nodes (MLN) were stimulated with PMA and Ionomycin in the presence of brefeldin A for 4 hr prior to intracellular cytokine staining (Panel E: 34-42 d post EC; separate mice from Panel B). Unstimulated cells received medium with only brefeldin A. The number of CD3+ CD4+ cells producing IL-17A or interferon-gamma is plotted. For ICC Isotype data points (iso), each sample was also stimulated with PMA/Ionomycin and stained for surface markers, with isotype control antibodies used for the intracellular cytokines. Line at median.Statistics: Mann-Whitney U test for all except Panel C, Wilcoxon matched-pairs signed rank test.

### 3.2 *E. coli* inoculum and enumeration

An overnight culture of *E. coli* SWW33 (ST375, [32]) was pelleted, resuspended in 20% glycerol, and stored at −80°C. 0.1 mL of the freshly thawed stock was administered by oral gavage. Fresh fecal pellets or colon contents collected immediately post-euthanasia were used for *E. coli* colony-forming unit (CFU) analysis. Approximately 5–10 mg of material was suspended in sterile phosphate-buffered saline (PBS) at a ratio of 10 mg per 1 mL of PBS and homogenized thoroughly by vortexing. Serial ten-fold dilutions were prepared in PBS, and aliquots were plated onto MacConkey II agar, which selectively supports *E. coli* growth under aerobic conditions while preventing the growth of ASF-associated bacteria. Plates were incubated at 37°C for ~15-24 hours and colonies were counted manually.

### 3.3 Altered Schaedler Flora (ASF)

ASF-colonized C3H mice (Model No. C3H) were purchased from Taconic Biosciences Inc (Germantown, NY). The germ-free shipper was attached to a proprietary port/cart unit, and the shipper/port was thoroughly sprayed with Spor-Klenz RTU. The port was fitted between the sash and the metal grate of the biosafety cabinet (BSC). The mice were sterilely transferred from the germ-free shipper into the BSC and placed into Allentown Sentry SPP™ (Sealed Positive Pressure) caging system (Allentown LLC, Allentown, PA). The mice were bred over a few months, and cecal contents were processed in an anaerobic chamber and stored at −80°C in sealed single-use glass vials. Each batch of inoculum was tested in germfree mice and confirmed to contain at least 7 of the 8 members (ASF360 is often below the limit of detection, or absent [25]). Mice used in studies were either born with ASF or colonized with ASF for at least 25d prior to administration of *E. coli*.

### 3.4 FITC-dextran assay

Intestinal barrier integrity was assessed using a fluorescein isothiocyanate–dextran (FITC-D) permeability assay [33]. Mice were fasted overnight (food only, water still present), followed by administration of 0.6 mg per 1 g of body weight by oral gavage. At 4 hr, mice were euthanized by carbon dioxide (CO_2_) asphyxiation, followed by cardiac puncture. Generally, ~0.5 mL of blood was collected via cardiac puncture and transferred to BD Microtainer® blood collection tubes. Samples were kept at room temperature in the dark for 30 minutes to allow for coagulation, then placed on ice, also in the dark, to preserve fluorescent signals and prevent degradation, followed by serum separation and storage at −80°C. Serum samples were diluted 1:1 with PBS and read on a Victor Nivo Multi-Mode Plate Reader with an excitation filter of 485 nm and an emission filter of 528 nm. A standard curve was generated using the same FITC-D preparation administered orally in that experiment.

### 3.5 Flow cytometry

Spleens and mesenteric lymph nodes (MLN) were mechanically dissociated into single-cell suspensions. Cells were stimulated for ~4 hours in medium with the presence of 10 µg/mL brefeldin A (Sigma-Aldrich, St. Louis, MO), 10 ng/mL Phorbol 12-myristate 13-acetate (Sigma-Aldrich), and 2.8 µM Ionomycin (Sigma-Aldrich. Live cell counts (obtained using a hemocytometer and trypan blue exclusion) were used to calculate a total cell count for each sample. After surface cell Fc receptors were blocked using Fc block (BD Biosciences, San Jose, CA), surface markers were stained with CD4 PerCP/Cyanine5.5, CD8 PE/Cyanine7, CD3ε Pacific Blue™ (all from Biolegend, San Diego, CA), and CD19 APC-Cy7 (BD Biosciences).

Cells were then fixed and permeabilized using BD Cytofix/Cytoperm™(BD Biosciences), and the intracellular cytokines were stained with IL-17A FITC, IL-2 APC (both from Biolegend), IFN-γ PE, and TNF Alexa Fluor® (both from BD Biosciences). Concurrently, isotype controls were stained with (rat) IgG1 FITC, IgG2b APC, IgG1 Alexa Fluor 700 (all from Biolegend), and IgG1 PE (BD Biosciences). After staining, cells were fixed in 1% paraformaldehyde, and data were acquired the following day using the BD LSRFortessa or BD FACSymphony A5 SE cell analyzers. Events were collected on high speed, recording 500,000 total events for spleen cells and 100,000 total events for mesenteric lymph node cells. Compensations were done after sample acquisition in FlowJo (FlowJo LLC, Ashland OR), using single stain controls that were stained and analyzed concurrently with the samples.

### 3.6 Pathology

Intestinal tissues from the proximal and distal small intestine were harvested with their contents, rolled into separate cassettes, fixed in 10% neutral buffered formalin, and routinely paraffin-embedded. Tissue sections (5 µm) were stained with hematoxylin and eosin and evaluated by a board-certified veterinary pathologist blinded to genotype and experimental manipulation. Slides were evaluated for the following parameters, all scored on a scale of 0-4: inflammation; mucosal proliferation (scored subjectively as increased mitoses, basophilia, piling of crypt epithelial cells, and relative elongation of the crypt to non-crypt villi above the expected baseline for that region of intestine); retention of Paneth cell contents, and mucus retention by goblet cells. In general, a score of 0 = normal; 1 = minimally increased; 2 = mildly increased; 3 = moderately increased; and 4 = severely increased. Images were captured from glass slides using NIS-Elements BR 4.20.01 and plated in Adobe Photoshop 2023. Image white balance, lighting, and contrast were adjusted using auto-corrections applied to the entire image. Original magnification is stated in Figure 4A.

### 3.7 16S rRNA sequencing

We performed 16S rRNA amplicon sequencing using methods previously published by our group [34–36]. Briefly, we extracted DNA from fecal pellets, using the Zymo Quick-DNA Fecal/Soil Microbe Miniprep Kit. For the first round of PCR, we used 10 – 25 ng input of extracted gDNA to prepare 25 uL reactions. We amplified the V4 hypervariable region of the16S rRNA gene using the locus specific sequences of the 515F & 806R primers per the earth microbiome project and Illumina’s forward and reverse overhang sequence. The forward and reverse primers were added to the reaction at 10 µM and Peptide Nucleic Acids (PNA) were added at 5 µM. The resulting PCR product was then used in library construction by ligation of Illumina’s unique dual indexes to the initial PCR product.

Quality control steps were performed on the extracted DNA before amplification of the V4 region of 16S rRNA gene and after library preparation. Each sample was quantified using the Qubit Assay Kit. The extracted gDNA and the resulting PCR products from library construction were verified using agarose gels. After the libraries were prepared, they were pooled to ensure equal sample distribution amongst the reads. Finally, the library pool was quantified using qPCR, this quantifies the concentration of library molecules that can be sequenced. Included in the final library pool were negative controls for DNA Extraction, PCR-1 (amplification of the v4 region) and PCR-2 (library construction). We also included ZymoBIOMICS Microbial Community Standard as a positive control. The final pool was sequenced on an Illumina sequencer. The sequencing parameters were single read 150 cycle.

We demultiplexed raw sequencing data using bcl2fastq v2.20 (Illumina) with default settings. First, we trimmed adapters and low-quality bases (Phred score < 20) using Trimmomatic v0.402. We removed chimeric sequences using the DADA2 consensus method [37]. We assigned taxonomy using the SILVA database (v138.1) with a minimum bootstrap confidence of 80 %. Downstream analysis of 16S rRNA amplicon data was performed using the phyloseq package in R [38].

Using vsearch (v0.1) [39] with a minimum global identity cut-off of 97%, 16S rRNA gene sequences were mapped to a custom database containing full-length 16S rRNA gene sequences for each ASF type strain and *E. coli* strain SWW33. Reads for each 16S rRNA gene within a species were pooled to obtain total reads per species. We used the Kruskal-Wallis test, corrected using the two-stage linear set-up procedure of Benjamini, Krieger, and Yekutieli in Prism software, to determine significance (q<0.05).

### 3.8 Dual RNA sequencing

Sections (3 – 4 cm) of distal small intestine from mice that were colonized with ASF + *E. coli* SWW33 were used to extract the total RNA using Qiagen’s RNeasy PowerFecal Pro kit. During RNA extraction all samples underwent Dnase treatment. Dnase treated total RNA samples were quantified using Qubit Assay kit and RNA quality was determined using Agilent’s Tapestation. We used the MicrobEnrich kit from Thermofisher to deplete mammalian RNA from each sample, thereby enriching bacterial RNA. Another round of QC was performed to ensure the integrity of the total RNA was not compromised during the enrichment process. The remaining volume of un-enriched total RNA and the enriched RNA were then used for library construction. Libraries were prepared using the Illumina Stranded Total RNA Ligation Prep with Rib-Zero Plus kit. The resulting library was quantified using the Qubit Assay kit and library quality was verified using Agilent’s TapeStation. The libraries were pooled at equimolar concentrations, and the library pool was quantified using qPCR to determine concentration of library molecules that can be sequenced. The library pool was sequenced on a NovaSeq X instrument. The sequencing parameters used were paired-end 150 cycles on 1 lane of a 10 billion-read flowcell.

We assessed raw FASTQ quality using FastQC v0.11.9. We trimmed adapters and low-quality bases (Phred < 20) using Trimmomatic v0.40 [40] with the following parameters: ILLUMINACLIP:QIAseq_adapters.fa:2:30:10, LEADING:20, TRAILING:20,

SLIDINGWINDOW:4:20, MINLEN:50. We removed contaminant host reads by mapping against the reference genome (GRCm39) using Bowtie2 v2.4.4 with --very-sensitive-local settings and discarding aligned reads [41]. We performed taxonomic classification of cleaned reads using Kraken2 v2.1.2 [42] with a custom database comprising RefSeq bacterial, archaeal, viral, and fungal genomes. We set the confidence threshold to 0.1. We generated per-sample taxonomic abundance reports using Bracken v2.7 [43] with a read length of 150 bp and k-mer distribution recalculated for our database. We summarized results at genus and species levels and converted raw counts to relative abundances.

### 3.9 Bioinformatics Analysis

Raw reads from host bulk-RNASeq data were quality-trimmed and adapter-removed using Trimmomatic (version 0.39) with default parameters for paired-end reads. Trimmed reads were then aligned to the mouse reference genome (GRCm39) using the STAR aligner (version 2.7.10a [44]) in two-pass mode to generate BAM files. Gene abundance matrices were created by quantifying aligned reads against the Ensembl gene annotation using featureCounts from the Subread package (version 2.0.3) [45]. Differential expression analysis was performed using DESeq2 (version 1.34.0) in R [46], followed by gene set enrichment analysis (GSEA) for pathway enrichment using the fgsea package (version 1.20.0) [47] with the KEGG database as the reference, applying a significance threshold of adjusted p-value < 0.05.

## 4. Results

### 4.1 Lack of disruption of intestinal epithelial barrier in germ-free (GF) or ASF-colonized CF mice

It is well established that conventional SPF mice with CF, as well as people with CF, show increased intestinal permeability [2–5]. We confirmed that we could measure this increased intestinal permeability using the FITC-dextran permeability assay in SPF CF mice (full-KO line G542X [31]) (**Figure 1A**). To test the role of the microbiota in intestinal barrier dysfunction, we performed the same assay in germ-free mice and found that the GF CF mice (original full-KO line S489X [29]) did not show an increase in permeability compared to control non-CF mice (**Figure 1A**). To determine whether proteobacteria contribute to barrier disruption, we next tested the defined 8-member bacterial community that lacks proteobacteria, Altered Schaedler Flora (ASF) [25], and found that ASF-colonized CF mice were no more permeable to FITC-dextran than controls (**Figure 1A**). This demonstrated that, in the absence of *E. coli*, CF mice colonized with ASF did not show increased intestinal permeability.

### 4.2 Addition of *E. coli* to ASF-colonized mice results in increased intestinal permeability and *E. coli* CFU in CF

We next orally inoculated ASF-colonized mice with a mouse commensal *E. coli* strain (SWW33/ST375, [32]). ASF does not mediate colonization resistance, allowing *E. coli* to colonize the mouse gut and be maintained long-term. We found that CF mice colonized with ASF + *E. coli* displayed significantly increased intestinal leakiness of FITC-D compared to control mice (**Figure 1A**). CF mice also had an increase in *E. coli* CFU in colon contents at necropsy compared to non-CF (**Figure 1B**).

### 4.3 TH17 responses are further enhanced in CF mice colonized with ASF and *E. coli* compared to ASF alone

Given our previous observation of increased TH17 lymphocytes in the original full-KO line S489X [22], we examined T helper cell differentiation in the ASF− and ASF+ *E. coli*-colonized mice (**Figure 1C,1D,1E**). We stimulated single-cell suspensions of whole splenocytes and mesenteric lymph nodes (MLN) with phorbol 12-myristate 13-acetate (PMA) and ionomycin and then stained for intracellular cytokines in CD4+ T cells. We hypothesized that increased intestinal epithelium permeability would drive an increase in TH17 cells, and predicted that CF mice colonized with only ASF, that did not display an increase in intestinal permeability (**Figure 1A**), would show limited TH17 cells. However, we found that ASF-colonized CF mice maintain higher numbers of TH17 cells in the spleen and MLN compared to control non-CF mice (**Figure 1C,1D**). To confirm that the staining was valid, we stimulated several samples with PMA/Ionomycin, stained them for surface markers, and used isotype control antibodies for intracellular cytokines (iso, in **Figure 1D**), which were negative for signal. Similarly to our previously published results [22], we also observed that T cells were already producing IL-17A prior to restimulation *in vitro* (see Unstimulated cells in MLN **Figure 1C,1D**). Note that the number of TH1 lymphocytes was equivalent between CF and non-CF mice (**Figure 1D**, bottom row), demonstrating that it is not a general pan-T-cell effect and that only TH17 cells are increased in CF. TH17 cells were further amplified in CF mice colonized with ASF + *E. coli* (**Figure 1C,1E**; median of 0.75 x 10e4 in ASF+EC, compared to 0.22 x 10e4 in ASF alone, **Figure 1C,1E**, in PMA/Ionomycin restimulated MLN).

### 4.4 Loss of *Cftr* in the intestinal epithelium is sufficient to phenocopy the full KO

To specifically examine the role of the intestinal epithelium in barrier permeability and *E. coli* expansion, we rederived the *Cftr*^*fl*/fl^ x Villin-Cre line where Cftr is only deleted in the intestinal epithelium by the use of a floxed *Cftr* allele (*Cftr*^*fl*/fl^) and Villin promoter-driven Cre recombinase expression [28]. Similar to the full KO mice, the intestinal KO (intKO) mice (i.e. *Cftr*^*fl*/fl^ Vil:Cre^+^) also showed increased intestinal permeability (**Figure 2A**) and increased colonic *E. coli* CFU (**Figure 2B**) compared to intestinal controls (intCtrl: *Cftr*^*fl*/fl^ Vil:Cre^-^ or *Cftr*^*fl*/+^ Vil:Cre^+or-^). Consistent with higher *E. coli* CFU and increased intestinal permeability, we also measured higher TH17 cells in the MLN (**Figure 2C,2D**), but surprisingly this increase was not observed in the spleen as seen with the full KO line (**Figure 1E**).

**Figure 2:**
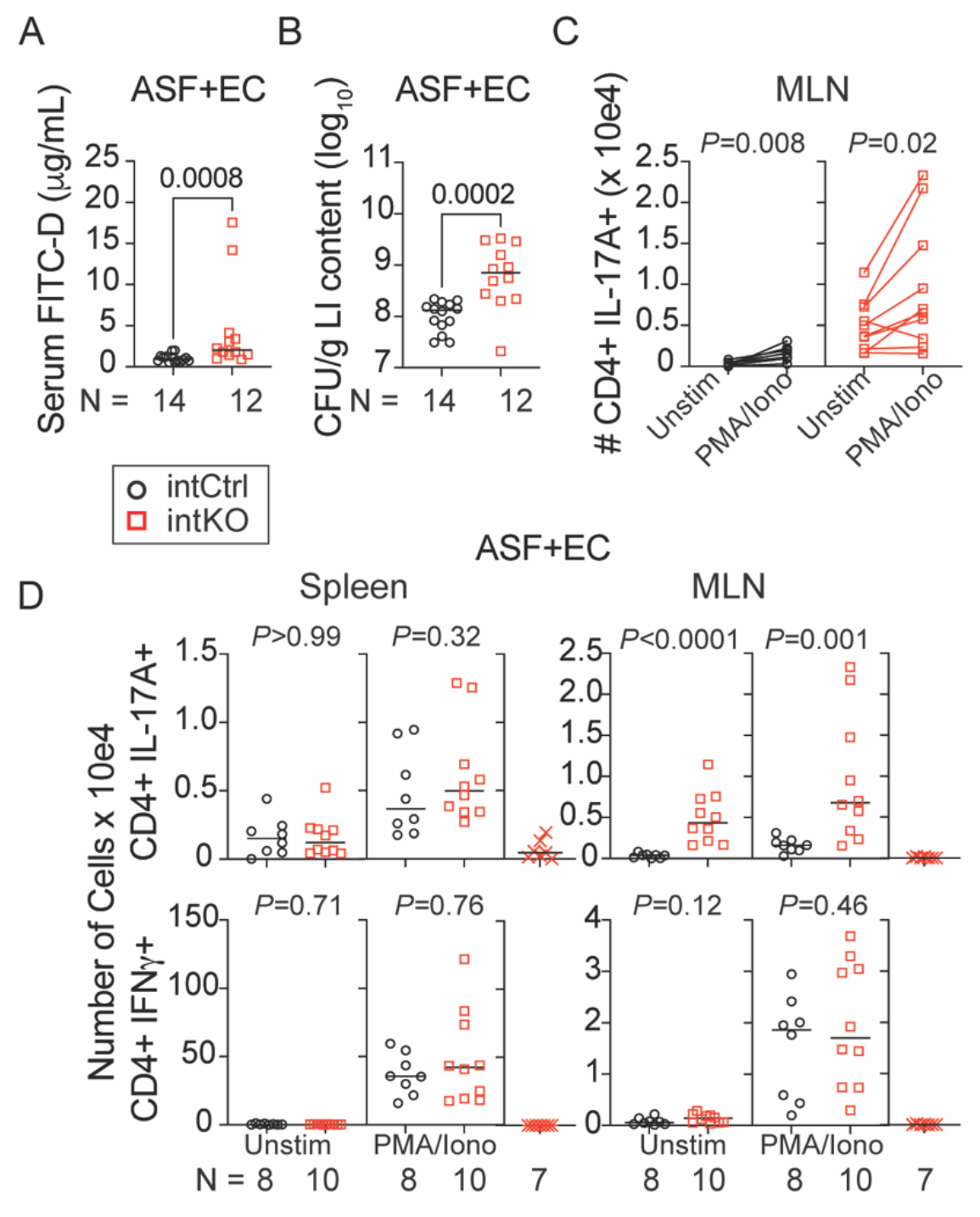
Mouse commensal *E. coli* strain SWW33 disrupts intestinal barrier in intKO mice. **Panel A)** 14-46 days after *E. coli* administration, FITC-Dextran was measured in the serum 4 hr after oral gavage. Note that the results in Panel A remain significant if the 2 high values are excluded from the intKO group (*P*=0.004). **Panel B)** Colon (LI) contents collected at necropsy were plated on MacConkey II Agar for *E. coli* CFU enumeration (same mice as Panel A). **Panels C & D)** 31-56 days after *E. coli* administration (different mice from Panels A & B), cells from spleens or MLN were stimulated with PMA and Ionomycin in the presence of brefeldin A for 4 hr prior to intracellular cytokine staining as in Figure 1. Line at median. Statistics: Mann-Whitney U test for all except Panel C, Wilcoxon matched-pairs signed rank test.

### 4.5 The proportion of *Eubacterium plexicaudatum* is significantly reduced in feces of CF and intKO mice

Analysis of ASF community structure in CF mice and controls via 16S rRNA gene (V4 region) profiling reveals that the proportion of ASF492 (*Eubacterium plexicaudatum*) is significantly reduced in CF mice both pre- and post-*E. coli* administration (**Figure 3**, left and middle panels). While there is a trend towards enrichment of ASF361 (*Ligilactobacillus murinus*) and a depletion of ASF519 (*Parabacteroides goldsteinii*) in CF mice prior to *E. coli* administration, the large spread of the data resulted in a lack of overall significance. However, these data suggest that CF can drive alterations in community structure even within a small defined community. Consistent with our CFU data, *E. coli* was found to be enriched in CF mice whereas ASF502 (*Schaedlerella arabinosiphila*) was depleted. Note that *E. coli* is the only bacterium whose absolute abundance (via CFU) was determined. Consistent with our results in the full KO line, ASF492 was also depleted in the intestinal KO mice (**Figure 3**, right panel). This organism was first isolated from healthy laboratory and wild mice in 1974 due to its distinctive subpolar tuft of flagella [49]. At the time, its production of high levels of butyric acid was proposed to inhibit expansion of coliform bacteria (including *E. coli*) in the cecum. Thus, its depletion in CF/intKO mice with increased *E. coli* growth is consistent with this hypothesis. However, we found higher levels of butyric acid in the feces of CF mice colonized with ASF + *E. coli* compared to control non-CF mice (data from one experiment with N=5/genotype; median of 0.21 umol/g feces in CF versus 0.06 umol/g feces in non-CF; not shown).

**Figure 3:**
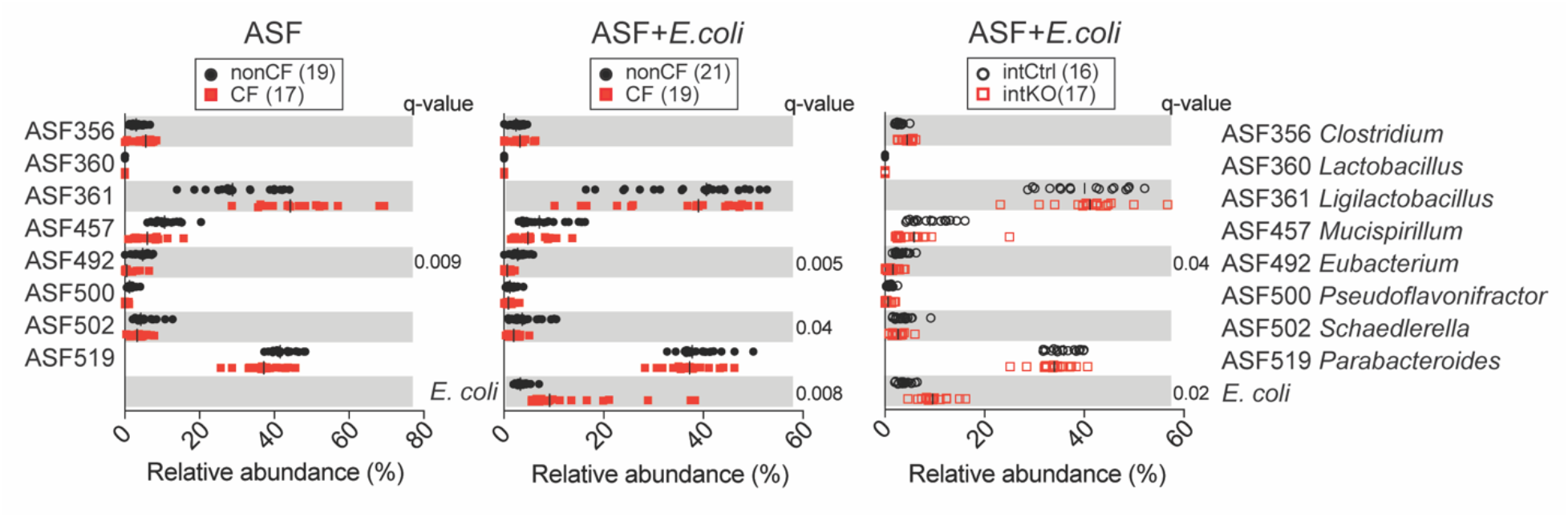
ASF community structure determined via 16S rRNA gene sequencing. The relative abundance of each ASF member (and *E. coli*, if present) is shown in fecal samples from CF and control non-CF mice (left and middle panels), or from intKO and respective intCtrl (right panel). Time of *E. coli* colonization ranged from 6 to 36 days for CF line, and 6 to 79 days for intKO line. Note that only a subset of the mice shown in the left and middle panels were assayed before and after *E. coli* administration. Statistics: Kruskal-Wallis followed by the 2-stage linear set-up procedure of Benjamini, Krieger and Yekutieli.

### 4.6 Similar histological changes in the distal small intestine of CF and intKO mice colonized with ASF + *E. coli*

We (JMS, veterinary pathologist blinded to genotype) examined the distal small intestine, the site of previously described histological changes in germ-free and SPF CF mice [13,22,29], of CF and intKO mice and their respective controls after colonization with ASF + *E. coli* (**Figure 4**). The CF and intKO mice displayed increased retention of Paneth cell contents in the crypts as well as mucus retention by goblet cells. All groups had increased crypt mitoses, potentially due to laxative treatment, although the CF and intKO mice had a subjectively increased crypt-to-non-crypt ratio of the villi, particularly in the distal intestine. Inflammation was minimal to mild (**Figure 4**). Thus, limiting deletion of *Cftr* to only the intestinal epithelial cells (i.e. the intKO) did not limit the pathology observed in the intestines compared to the whole-body KO (i.e., CF mice, **Figure 4B**).

**Figure 4:**
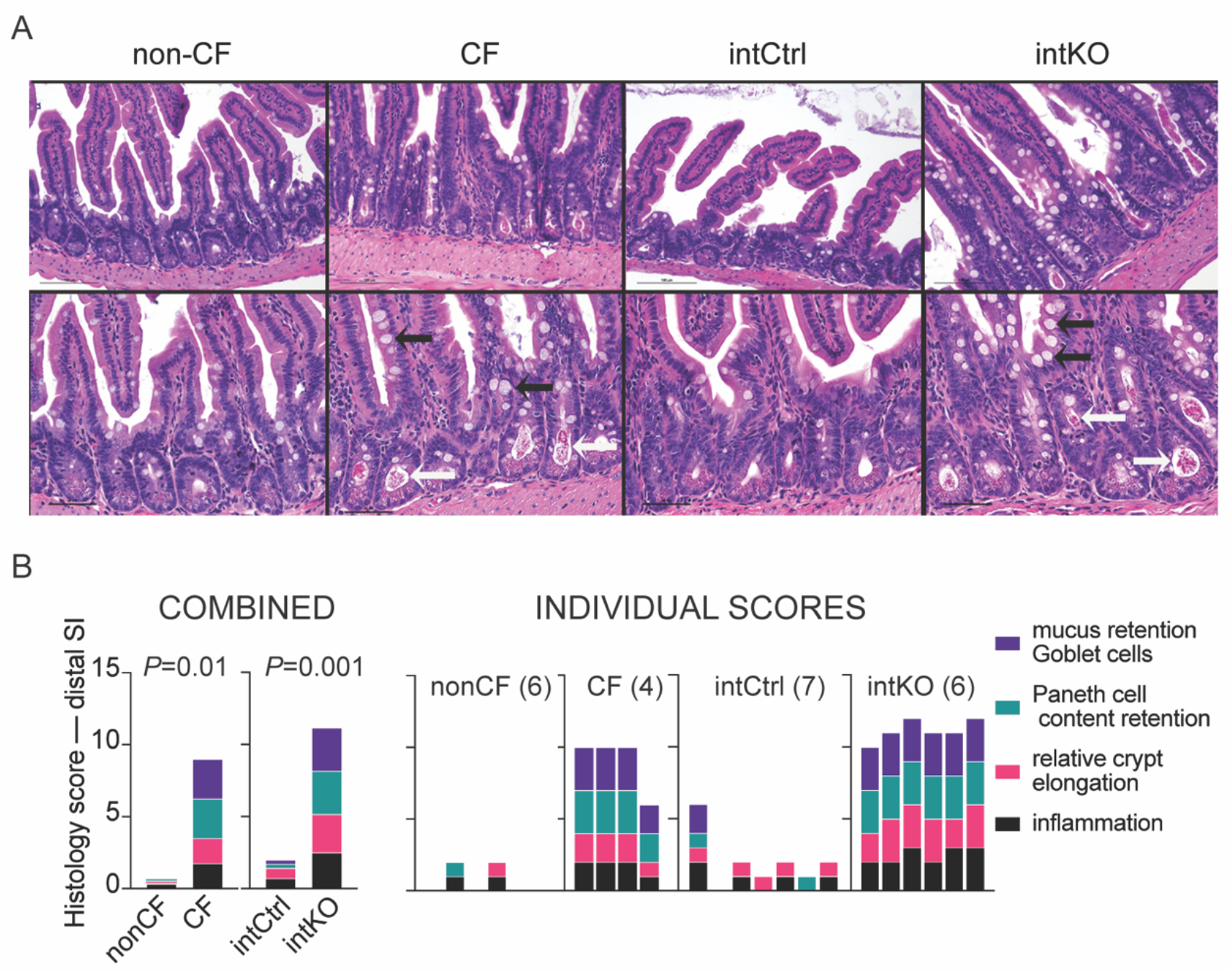
Histological analysis of intestinal sections of mice colonized with ASF + *E. coli*. **Panel A)** H&E staining of distal small intestine (SI). Left to right: non-CF (control); CF (KO); intCtrl (*Cftr*^*fl*/fl^ Vil:Cre-); intKO (*Cftr*^*fl*/fl^ Vil:Cre+). Top row: 200X original magnification, bar = 100 microns. Bottom row: 400X original magnification, bar = 50 microns. White arrows: retention of Paneth cell contents in the crypts. Black arrows: mucus retention by goblet cells. **Panel B)** Combined scores of histology by genotype on Left side (means of each feature), with individual scores for each mouse shown on Right side. Statistics: pairwise Mann-Whitney on cumulative total scores for each line of mice.

### 4.7 Common host pathways enriched and depleted in distal intestinal sections of CF and intKO mice following colonization with ASF + *E. coli*

To examine global host responses to colonization with ASF + *E. coli*, we extracted total RNA from whole sections (including the digesta) of distal small intestine from CF and intKO mice and their respective controls (the same mice from Figure 4 were used for these analyses). Bulk RNA sequencing was performed on the distal small intestine and enriched or depleted host KEGG pathways are shown in **Figure 5A,5B**. Several pathways were conserved between CF and intKO mice (the names are color coordinated in the figure) demonstrating that loss of *Cftr* in just the intestinal epithelium is sufficient to trigger CF-related changes in the gut.

**Figure 5:**
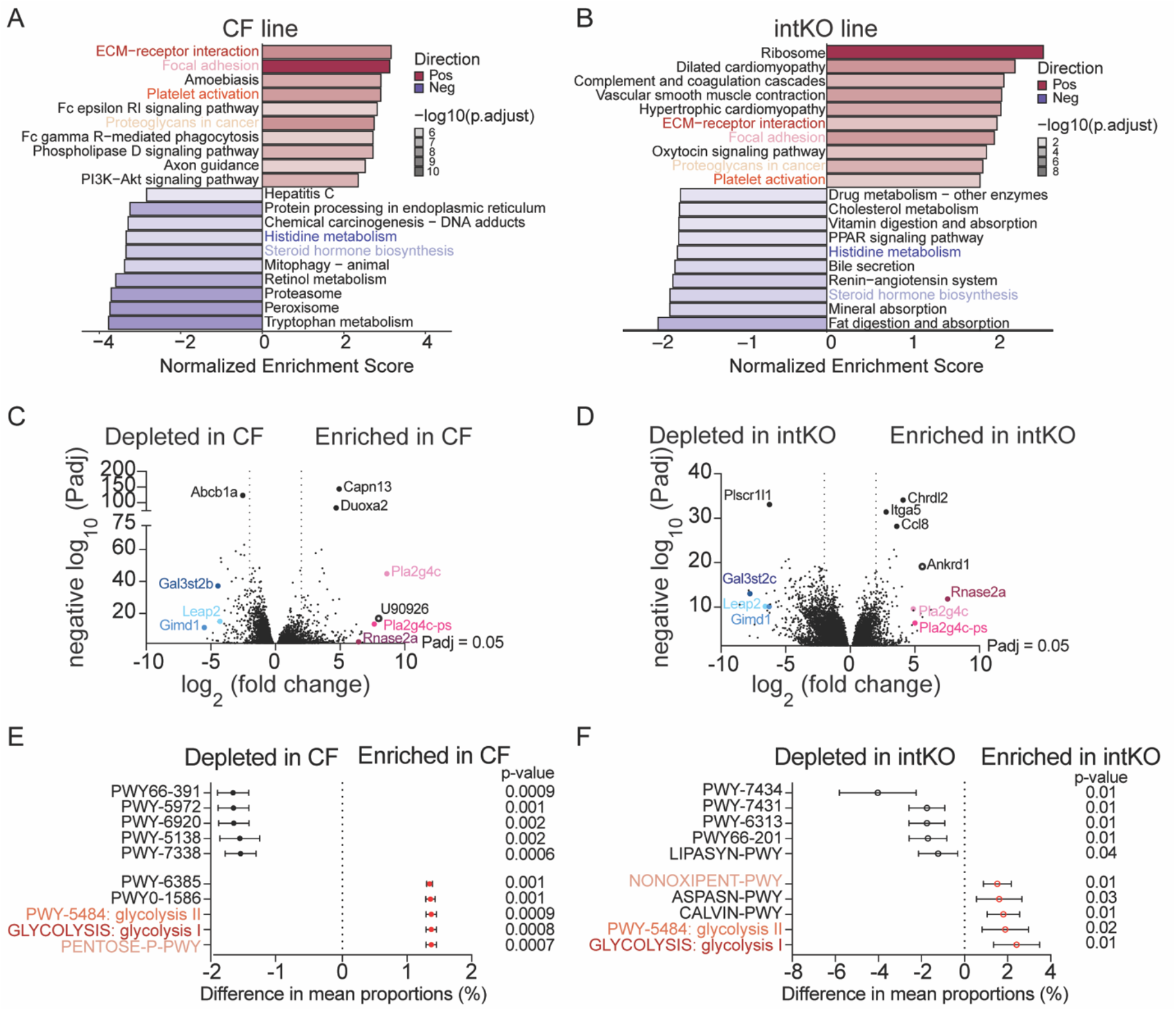
Shared host and bacterial pathways / genes in dSI colonized with ASF + *E. coli*. **Panels A & B)** Bi-directional bar plot illustrating the significant enrichment of KEGG pathways identified through gene set enrichment analysis (GSEA) from the same mice as in Figure 4. CF line: CF = 4, non-CF = 6 (Panel A). intKO line: intKO = 6, intCtrl = 7 (Panel B). The y-axis lists the pathway names, while the x-axis represents the Normalized Enrichment Score (NES). Bars extending to the right (positive NES) indicate pathways significantly upregulated in the KO or intKO group compared to the respective control group, whereas bars extending to the left (negative NES) indicate downregulated pathways. The bar color corresponds to the adjusted p-value (or FDR), with darker shades indicating greater statistical significance. The bar length reflects the magnitude of the enrichment score. The names of the pathways shared by CF and intKO lines are highlighted. Only pathways with an adjusted p-value < 0.05 are included. **Panels C & D)** Volcano plots depicting all significantly (P_adj_ <0.05) up and down regulated genes in CF (**Panel C**) and intKO (**Panel D**) mice compared to the respective controls. Note that the shared genes shown in Table 1 are highlighted in color in both panels. Dotted vertical lines depict cut-off for 4-fold up or down regulated genes. **Panels E & F)** Difference in mean proportions, with 95% Confidence Interval error bars, is plotted for the top 5 bacterial pathways enriched or depleted in CF (**Panel E**) or intKO (**Panel F**) mice. Corrected p-values are shown for each pathway. Shared pathways are highlighted. PWY66-391: fatty acid β-oxidation VI (mammalian peroxisome); PWY-5972: stearate biosynthesis I (animals); PWY-6920: 6-gingerol analog biosynthesis (engineered); PWY-5138: fatty acid β-oxidation IV (unsaturated, even number); PWY-7338: 10-trans-heptadecenoyl-CoA degradation (reductase-dependent, yeast); PWY-6385: peptidoglycan biosynthesis III (mycobacteria); PWY0-1586: peptidoglycan maturation (meso-diaminopimelate containing); PWY-5484: glycolysis II (from fructose 6-phosphate); GLYCOLYSIS: glycolysis I (from glucose 6-phosphate); PENTOSE-P-PWY: pentose phosphate pathway; PWY-7434: terminal O-glycans residues modification (via type 2 precursor disaccharide); PWY-7431: aromatic biogenic amine degradation (bacteria); PWY-6313: serotonin degradation; PWY66-201: nicotine degradation IV; LIPASYN-PWY: phospholipases; NONOXIPENT-PWY: pentose phosphate pathway (non-oxidative branch) I; ASPASN-PWY: superpathway of L-aspartate and L-asparagine biosynthesis; CALVIN-PWY: Calvin-Benson-Bassham cycle.

We examined the top 10 most upregulated or downregulated genes and identified shared genes between both CF and intKO mice (**Table 1** and **Figure 5C,5D**). Loss of *Cftr* in the intestinal epithelium was sufficient to result in strong upregulation of phospholipase A2 group IVC (*Pla2g4c*, and its pseudogene *Pla2g4c-ps*) as also seen in the whole-body KO CF line. This gene is found in the Focal Adhesion and Platelet Activation KEGG pathways that are upregulated in both lines (**Figure 5A,5B**). Upregulation of *Pla2g4c* (as well as phospholipase A2, group V) was previously demonstrated in SPF CF pups starting at weaning (P24) and increasing into adulthood [50]. *Rnase2a*, an eosinophil-associated RNase (EAR) gene in the RNase A superfamily, is highly upregulated in both lines and the human gene *Rnase2* (also known as eosinophil-derived neurotoxin, EDN [51]) is found in the shared Proteoglycans in Cancer KEGG pathway. The highly upregulated long noncoding RNA *U90926* is not detected in the intKO line but is the second most highly induced gene in CF mice colonized with ASF + *E. coli* {log_2_(FC) = 7.98, P_adj_ =1.47E-17}. In terms of innate immunity, *U90926* has been shown to be induced by toll-like receptor signaling and to encode a secreted protein that protects against endotoxic shock [52]. The other highly induced gene that appears in the intKO, Ankyrin repeat domain 1 (*Ankrd1*, **Figure 5D**), is also detected in CF, but not in the top 10 {log_2_(FC) = 4.48, P_adj_ = 0.0035}, and is part of Focal Adhesion, and Proteoglycans in Cancer KEGG pathways.

**Table 1:** Genes that exhibit the highest up- or down-regulation (by fold-change) shared by both CF and intKO mice.

| Gene | CF line |  | intKO line |  |
| --- | --- | --- | --- | --- |
| | $\log_2(\text{FC}^*)$ | Padj | $\log_2(\text{FC})$ | Padj |
| *Pla2g4c* | 8.6 | 1.4E-45 | 4.9 | 2.5E-10 |
| *Pla2g4c-ps* | 7.6 | 4.0E-14 | 5.0 | 4.0E-07 |
| *Rnase2a* | 6.4 | 0.006 | 7.5 | 1.5E-12 |
| *Gimd1* | -5.5 | 6.0E-12 | -6.3 | 7.8E-11 |
| *Gal3st2b* | -4.5 | 5.3E-38 |  |  |
| *Gal3st2c* |  |  | -7.8 | 9.8E-14 |
| *Leap2* | -4.3 | 8.0E-16 | -6.6 | 6.8E-11 |

\*FC = fold change

Genes with the most significant P_adj_ scores are also called out in **Figure 5C,5D**. Genes upregulated in CF mice include *Capn13* (encodes Calpain-13, a calcium-dependent cysteine endopeptidase) that is found in the Proteoglycans in Cancer pathway, similar to *Duoxa2* (dual oxidase maturation factor 2) that is additionally found in the ECM-Receptor Interaction and Platelet Activation pathways. Note that both Capn13 and Duoxa2 are also significantly upregulated in the intKO line, but the P_adj_ values are much lower (negative log_10_ of ~4-5). For the intKO, *Itga5* (integrin alpha-5) is found in all 4 shared upregulated pathways (ECM-Receptor Interaction, Focal Adhesion, Platelet Activation, and Proteoglycans in Cancer), whereas *Chrdl2* (Chordin-like 2) and *Ccl8* (C-C motif chemokine ligand 8) are not in any of the shared pathways. Both *Itga5* and *Ccl8* are below our log_2_FC cutoff of 2 in the CF line (and a negative log_10_ of <3), whereas *Chrdl2* (log_2_FC of 4.9) has a negative log_10_ of 3.1. None of the top downregulated genes are found in the shared KEGG pathways shown in **Figure 5A,5B**. Altogether, these results suggest that shared pathways and genes are induced in both CF and intKO mice colonized with ASF and *E. coli*, demonstrating the crucial role that CFTR plays in intestinal epithelial cells.

From the same distal intestinal samples, we also examined bacterial RNA expression and found shared pathways enriched in the microbiome of CF and intKO mice relative to their respective controls (**Figure 5E,5F**). Glycolysis and the pentose phosphate pathway are upregulated in both the CF and intKO mice, demonstrating enhanced bacterial metabolic activity when *Cftr* is mutated in the intestines, consistent with the higher CFU observed in both lines.

## 5. Discussion

Our results demonstrate that the intestinal epithelium is the key driver of microbiome changes in the intestinal tract in cystic fibrosis. Of note, this result demonstrates that the pancreatic insufficiency of CF and its resultant intestinal malabsorption is not necessary for the dysbiosis observed in CF individuals. Germ-free CF mice display entrapment of Paneth cell contents in dilated crypts in the distal small intestine [22] that undoubtedly alters selective pressure on the microbiota allowing for expansion or depletion of differential members of the microbial community living in the gut. This selective pressure is demonstrated even with the small defined ASF community, where depletion of ASF492 is observed in CF mice colonized with only ASF (Figure 3). Upon additional colonization with *E. coli*, we find that *E. coli* expands to higher numbers (both relative and absolute) in CF and intKO mice. *E. coli* is a facultative anaerobe and hence can survive in oxygenated tissue unlike members of ASF, aside from *Lactobacillus*, that are anaerobes. In addition, commensal *E. coli* strains can survive in close proximity to the epithelium even in the presence of inflammation by using nitrate as an electron acceptor during anaerobic respiration [53]. The unique environment of the CF gut allows *E. coli* to expand through a mechanism driven by the intestinal epithelium that is yet to be defined.

One advantage of studying the microbiome in mice is that we can use a defined microbiota and attribute changes solely to the genotype of the mouse. This is because control mice are housed under identical conditions, with identical water and food, excluding these factors from driving the observed changes. We previously showed that germ-free CF mice had a small but measurable population of TH17 cells in the MLN, despite lacking a microbiota. Here, we find that CF mice colonized with ASF have an increase in TH17 cells in both the spleen and the MLN, and that even in the absence of *in vitro* restimulation, the MLN harbors CD4+ cells producing IL-17A (Figure 1C and 1D). Upon colonization with *E. coli*, TH17 cells were further increased in the MLN (Figure 1C and 1D). Furthermore, an increase in intestinal permeability was observed in CF mice colonized with ASF + *E. coli*, that was absent in germ-free or ASF-colonized mice without *E. coli* (Figure 1A).

Surprisingly, despite increased intestinal permeability in the intKO mice, as well as increased *E. coli* CFU in the colon content, we did not observe increased systemic TH17 cells in the spleen (Figure 2). Whether this is due to lower levels of FITC-dextran measured in the serum compared to the full KO or due to lower *E. coli* expansion is unknown at this time. Global analysis of host responses provides a clue. The *U90926* gene, that protects against endotoxic shock, is the second most highly enriched gene in CF mice but is undetected in intKO mice.

This suggests that greater *E. coli* expansion and increased barrier disruption lead to greater innate immune stimulation. Furthermore, these results provide a rationale for reducing *E. coli* and other proteobacteria that expand in the CF gut.

In these studies, we used a commensal *E. coli* strain (SWW33) that had been found to promote higher levels of intestinal inflammation via differential interleukin-6 production in DSS-treated wild-type mice colonized with ASF [32]. Future studies will determine whether only a subset of commensal *E. coli* strains expand in CF and disrupt barrier function, or whether all strains share this function, e.g., due to LPS recognition by the innate immune response.

## 6. Acknowledgments

We thank the Cleveland Clinic Research Gnotobiotics Facility, Microbial Sequencing and Analytics Resource, Flow Cytometry Core and Histology Core. Funding for this work was provided by the Cystic Fibrosis Foundation.

## 7. Author Contributions

Study concept and design: PPA, NS, AMO, SIM, JM, AMH; execution of experiments: PPA, NS, JMS, ZJ, MC, BC, AM, LS, FS, BB, AkS, TN, CH, AMH; data analysis: PPA, NS, ApS, JMS, AMH; material support: MW, MD, MLD, CH; drafting of manuscript: PPA, NS, ApS, JMS, ZJ, MC, BB, TN, MD, MLD, CH, AMH; funding acquisition: AMH. All authors approved the manuscript.

## Notes

### Competing Interest Statement

The authors have declared no competing interest.

